# *eyes absent* in the cockroach panoistic ovaries regulates proliferation and differentiation through ecdysone signalling

**DOI:** 10.1101/839639

**Authors:** S. Ramos, F. Chelemen, V. Pagone, N. Elshaer, P. Irles, M.D. Piulachs

## Abstract

Eyes absent (Eya), is a protein structurally conserved from hydrozoans to humans, for which two basic roles have been reported: it can act as a transcription cofactor and as a protein tyrosine phosphatase. Eya was discovered in the fly *Drosophila melanogaster* in relation to its function in eye development, and the same function was later reported in other insects. Eya is also involved in insect oogenesis, although studies in this sense are limited to *D. melanogaster*, which has meroistic ovaries, and where *eya* mutations abolish gonad formation.

In the present work we studied the function of *eya* in the panoistic ovary of the cockroach *Blattella germanica*. We show that *eya* is essential for correct development of panoistic ovaries. In *B. germanica, eya* acts at different level and in a distinct way in the germarium and the vitellarium. In the germarium, *eya* contributes to maintain the correct number of somatic and germinal cells by regulating the expression of steroidogenic genes in the ovary. In the vitellarium, *eya* facilitates follicle cells proliferation and contributes to regulate the cell program, in the context of basal ovarian follicle maturation. Thus, *eya*-depleted females of *B. germanica* arrest the growth and maturation of basal ovarian follicles and become sterile.

## 1. Introduction

Maintaining the stability of stem cells is crucial in every organism, and this is especially important in the case of germinal stem cells. Oogenesis entails the process of ovary development from the time of germinal stem cell differentiation until oocyte maturation. During the oogenesis of many species, oocytes support a high level of transcription that is crucial not only for the growth of the oocyte, but also for the zygote activation, thus ensuring successful reproduction (Song and Wessel, 2005). In insects with meroistic ovaries, the transcriptional activity is mainly executed by the specialized nurse cells. Conversely, in panoistic ovaries the germinal vesicle is the responsible to maintain the oocyte itself, and provide the necessary materials to ensure embryogenesis (Bogolyubov, 2007; Büning, 1994).

In insect ovaries, germinal stem cells are located in niches in the germarium of each ovariole. The control of their proliferation and differentiation has been thoroughly studied in species with meroistic polytrophic ovaries, such as the fruit fly *Drosophila melanogaster* (see Ameku et al., 2017; Belles and Piulachs, 2015; Dai et al., 2017). In contrast, the knowledge of genes involved in regulating oogenesis in panoistic ovaries is very limited. Oocyte growth and maturation in panoistic ovaries has been systematically studied in the cockroach *Blattella germanica* (Elshaer and Piulachs, 2015; Herraiz et al., 2014; Irles et al., 2016; Irles and Piulachs, 2011; Tanaka and Piulachs, 2012), which is emerging as a choice model to study this ovary type. In *B. germanica*, each ovary has around 20 ovarioles, but only the most basal ovarian follicle of each ovariole matures during a given gonadotrophic cycle. The basal ovarian follicles are almost ready to mature in freshly ecdysed females, which means that the first gonadotrophic cycle starts early in the last nymphal instar. Importantly, the development of the remaining ovarian follicles of each ovariole is arrested until these basal ones are oviposited (Irles and Piulachs, 2014).

In previous contributions dealing with the regulation of oogenesis in panoistic ovaries, we have studied the function of *Notch* (*N*) in the ovary of *B. germanica* and its interactions with the EGFR signalling pathway (Elshaer and Piulachs, 2015; Irles et al., 2016; Irles and Piulachs, 2014). Given that the Notch pathway participates in the control of germinal cell proliferation, here we postulated that the main effectors of this function would be downstream genes in the same pathway. Thus, we focused on *eyes absent* (*eya*) as a candidate to play a key role in this process.

The *eya* gene is structurally conserved from hydrozoans to humans (Duncan et al., 1997; Graziussi et al., 2012; Jemc and Rebay, 2007). It was discovered in *D. melanogaster* for its role in determining cell fates (differentiation or death) in postembryonic development in relation to eyes formation (Bonini et al., 1993). Subsequently, *eya* orthologues have been found in vertebrates and in other phyla (Duncan et al., 1997; Graziussi et al., 2012; Zimmerman et al., 1997), and a wide range of functions in development have been reported for the corresponding protein.

Two basic roles have been reported for Eya. It was first described as transcriptional cofactor that is recruited to transcriptional complexes via the Eya domain (ED), a conserved C-terminal motif that interacts with the Six family DNA binding proteins (Jemc and Rebay, 2007). Subsequently, different research groups reported that the ED has intrinsic protein tyrosine phosphatase activity, establishing Eya as an example of a new class of eukaryotic protein phosphatases (see Rebay, 2015 and references therein).

In insects, the function of *eya* in eye development has been reported both in holometabolan species like *D. melanogaster* and the red flour beetle *Tribolium castaneum* (Yang et al., 2009), as well as in hemimetabolan species. In the later, Dong and Friedrich (2010) studied *eya* in the post-embryonic development of the locust *Schistocerca americana* and Takagi and co-workers (2012) investigated its role in eye development in embryos and nymphs of the cricket *Gryllus bimaculatus*.

Research on *eya* functions in insect oogenesis has been limited to *D. melanogaster*, where it was studied during gonad formation in embryogenesis (Boyle et al., 1997), and in the adult during egg chambers differentiation (Bonini et al., 1998). From the early stages of ovarian development (stage 2a) *eya*, is expressed in follicle cells until stage 10 of egg chamber, when follicle cells start the migration over the oocyte. An inappropriate follicle cell development results in sterility due to the arrest of egg chamber development (Bonini et al., 1998). More recently it has also been demonstrated that *eya* expression in the *D. melanogaster* ovary is essential to regulate polar and stalk cell fates. Thus, loss of *eya* transforms epithelial follicle cells in polar cells, while repression of *eya* is required for stalk cell formation (Bai and Montell, 2002).

In contrast to the research carried out in *D. melanogaster*, the possible role of *eya* in other insect ovary types remains unstudied. Therefore, in this work we aimed at addressing the role of this important protein in panoistic ovaries. We used the cockroach *B. germanica* as a model, and focussed on the regulation of cell proliferation and differentiation in the germarium.

## 2. Material and Methods

### 2.1. Cockroach colony and sampling

Adult females of the cockroach *B. germanica* (L.) were obtained from a colony fed *ad libitum* on Panlab dog chow and water, and reared in the dark at 29 ± 1°C and 60–70% relative humidity. Freshly ecdysed adult females were selected and used at appropriate ages. Mated females were used in all experiments (the presence of spermatozoa in the spermatheca was assessed at the end of the experiment to confirm mating. All dissections and tissue samplings were performed on carbon dioxide-anaesthetized specimens.

### 2.2. RNA extraction and expression studies

Total RNA was isolated using the GenElute Mammalian Total RNA Kit (Sigma, Madrid, Spain). A total of 300 ng from each RNA extraction was treated with DNAse (Promega, Madison, WI, USA) and reverse transcribed with Superscript II reverse transcriptase (Invitrogen, Carlsbad CA, USA) and random hexamers (Promega). RNA quantity and quality were estimated by spectrophotometric absorption at 260/280 nm in a Nanodrop Spectrophotometer ND-1000® (NanoDrop Technologies, Wilmington, DE, USA).

The expression pattern of the examined *B. germanica* genes was determined by quantitative real time PCR (qRT-PCR) in ovaries from sixth instar nymph and adults. One ovary pair, for adults, or pools of two ovary pairs for nymphs, for every chosen age were used. The expression levels in treated individuals were quantified individually. PCR primers used in qRT-PCR expression studies were designed using the Primer3 v.0.4.0 (Rozen and Skaletsky, 2000). The *actin-5c* gene of *B. germanica* (Accession number AJ862721) was used as a reference for expression studies. qRT-PCR reactions were made using the iTaq Universal SYBR Green Supermix (BioRad) containing 200 nM of each specific primer (performed in triplicate). Amplification reactions were carried out at 95°C for 2 min, and 40 cycles of 95°C for 15 s and 60°C for 30 s, using MyIQ Single Color RTPCR Detection System (BioRad). After the amplification phase, levels of mRNA were calculated relative to *actin-5c*. Results are given as copies of mRNA per 1000 copies of *actin-5c* mRNA. The primer sequences used to quantify gene expression are indicated in Table S1.

### 2.3. RNAi experiments

To deplete the expression of *eya*, two dsRNA (ds*eya*) were designed targeting the C-terminal domain of *eya* (318 and 325 bp each). As the same ovary phenotype was found using both dsRNA, we will refer to the RNAi treatments as ds*eya*. A dsRNA (dsMock) corresponding to 307-bp of the *Autographa californica* nucleopoyhedrovirus sequence was used as control. The dsRNAs were synthesized in vitro as we previously described (Ciudad et al., 2006). The dose used was 1 μg for either ds*eya* or dsMock, and they were injected into the abdomen of 0-day-old sixth nymphal instar or in 0-day-old adult females.

### 2.4. 20-Hydroxyecdysone treatments

Newly emerged last instar nymphs or adult females, were injected with 1µL of a 10mM 20-hydroxyecdysone (20E) (10% ethanol). Nymphs were dissected when they were 6-day-old, just when ecdysteroids in the hemolymph reaches the highest levels (Cruz et al., 2003), or when they were 8-day-old, thus just before the molt to adult stage. Adult females were dissected when they were 5-day-old, before choriogenesis begins.

### 2.5. Immunohistochemistry

After dissection, ovaries were immediately fixed in paraformaldehyde (4 % in PBS) for 2 h. Washing samples and antibody incubations were performed as previously described (Irles and Piulachs, 2014). The primary antibody employed were rabbit antibody anti-PH3 (Cell Signaling Technology, Denver, MA; dilution 1:250), rabbit antibody anti-cleaved Caspase-3 (Asp-175, Cell Signaling Tech; dilution 1:50), and mouse antibody anti-Eya, (deposited to the DSHB by Benzer, S. / Bonini, N.M.; product eya10H6; dilution 1:50) as nuclear marker of germinal cells. However, were unable to assess Eya labelling, since there is not decrease of protein labelling in dsRNA treated insects. The secondary antibodies used were Alexa-Fluor 647 conjugated donkey anti-rabbit IgG, or Alexa-Fluor 647 conjugated goat anti-mouse IgG (Molecular Probes, Carlsbad, CA). Ovaries were also incubated at room temperature for 20 min in 300 ng/ml phalloidin-TRITC (Sigma) and then for 5 min in 1 µg/ml DAPI (Sigma) PBT, to show the F-actin and nuclei, respectively. After three washes with PBT, ovaries were mounted in Mowiol (Calbiochem, Madison, WI, USA) and observed using a Zeiss AxioImager Z1 microscope (Apotome) (Carl Zeiss MicroImaging).

The number of cells in the follicular epithelia was estimated applying the function described in (Pascual et al., 1992).

We considered that an ovarian follicle has been released from the germarium when it is possible to identify the cell membrane surrounding the oocyte. The most basal follicle was excluded when quantifying ovarian follicles in the vitellarium.

### 2.5. Statistics

Quantitative data are expressed as mean ± standard error of the mean (S.E.M.). Statistical differences between morphometric data were evaluated using the ANOVA or the Student’s t-test using IBM SPSS statistics software. Comparisons of gene expression between treatment and control groups were made using the Pair-Wise Fixed Reallocation Randomization Test (which makes no assumptions about distributions) (Pfaffl et al., 2002), employing REST 2008 v. 2.0.7 software (Corbett Research).

## 3. Results

### 3.1. *eya* in *Blattella germanica* ovaries and efficiency of RNAi treatments

In the *B. germanica* ovaries, *eya* is expressed through the gonadotropic cycle (Figure 1A). The expression is remarkably variable in the last nymphal instar, although a clear peak can be observed just after the imaginal molt. Then, the expression begins to decline and reaches the lowest values on day 6, when choriogenesis starts. This pattern suggests that *eya* plays important functions in the early steps of oogenesis. To test this hypothesis, we used RNAi approaches. Thus, newly emerged sixth instar female nymphs, were treated with ds*eya* (n = 36) or dsMock (n = 40). All the ds*eya* treated nymphs molted to the adult stage, which indicates that the treatment does not affect the image molt, but all resulting adult females failed to oviposit (Table S2). To assess the efficiency of the RNAi, we did new treatments of freshly emerged sixth instar female nymphs with ds*eya*, and transcript decrease was examined in the ovaries at different ages. At the end of the nymphal stage (8-day-old sixth instar nymphs), *eya* mRNA levels were depleted (p = 0.097), then, the expression kept decreasing in 3- and 5-day-old adult females (p = 0.031 and p = 0.0001, respectively, Figure 1B).

**Figure 1.**
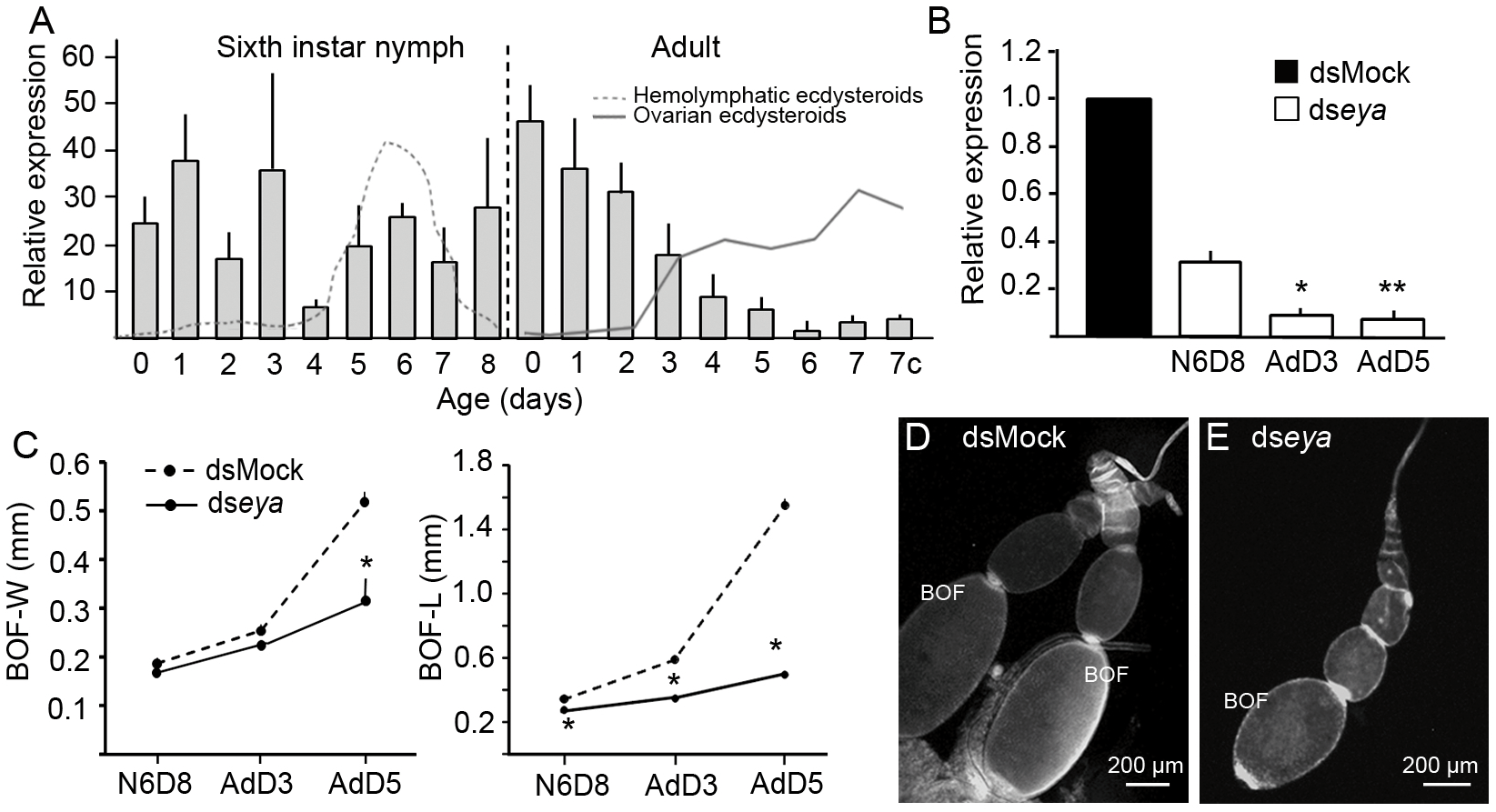
Expression of *eya* in the ovaries of *Blattella germanica* and effects of eya depletion. **A**. Expression pattern of *eya* in ovaries in sixth instar nymphs and adult females during the first gonadotropic cycle. Data represent copies of *eya* mRNA per 1000 copies of *actin-5c* and are expressed as the mean ± S.E.M. (n= 3-6). Profiles of ecdysteroids titer in the haemolymph (grey dashed line), and ecdysteroids content in the ovaries (grey solid line) are also shown (ecdysteroids data from Cruz et al., 2003; Pascual et al., 1992; Romaña et al., 1995). **B**. Expression of *eya* in ovaries of 8-day-old (N6D8) sixth instar nymph; 3-day-old adult (AdD3) and 5-day-old adult (AdD5) that were treated with ds*eya* in freshly emerged sixth nymphal instar. Data represent normalized values against dsMock-treated insects (reference value 1), and are expressed as the mean ± S.E.M. (n = 3). The asterisks indicate statistically significant differences with respect to controls: *, p = 0.031 and **, p= 0.0001). **C**. Width (BOF-W) and the length (BOF-L) of basal ovarian follicle in *eya*-depleted N6D8, AdD3 and AdD5 females. The asterisk indicates statistically significant differences with respect to controls: *, p < 0.0001). **D**. Ovarioles from N6D8 dsMock-treated female. **E**. Ovariole from an N6D8 ds*eya*-treated female.

### 3.2. *eya* is involved in growth and maturation of basal ovarian follicles

The growth of the basal ovarian follicles in *eya*-depleted females was slowed (Figure 1C), and their general shape became spherical (Figure 1D and E). There were significantly fewer follicular cells in the basal ovarian follicles of *eya*-depleted females compared to dsMock-treated females (Figure 2A). In 8-day-old sixth instar nymphs, labelling with PH3 antibody revealed fewer mitotic divisions in the follicular epithelia from *eya*-depleted females (Figure 2B and C), and the follicular cells within the epithelia in basal ovarian follicles showed a remarkable variation of nuclei size (Figure 2D and F). Additionally, F-actin appeared to concentrate at the junctions between follicular cell membranes, which could explain the changes in cell shape, and distribution through the epithelia (Figure 2E and G).

**Figure 2.**
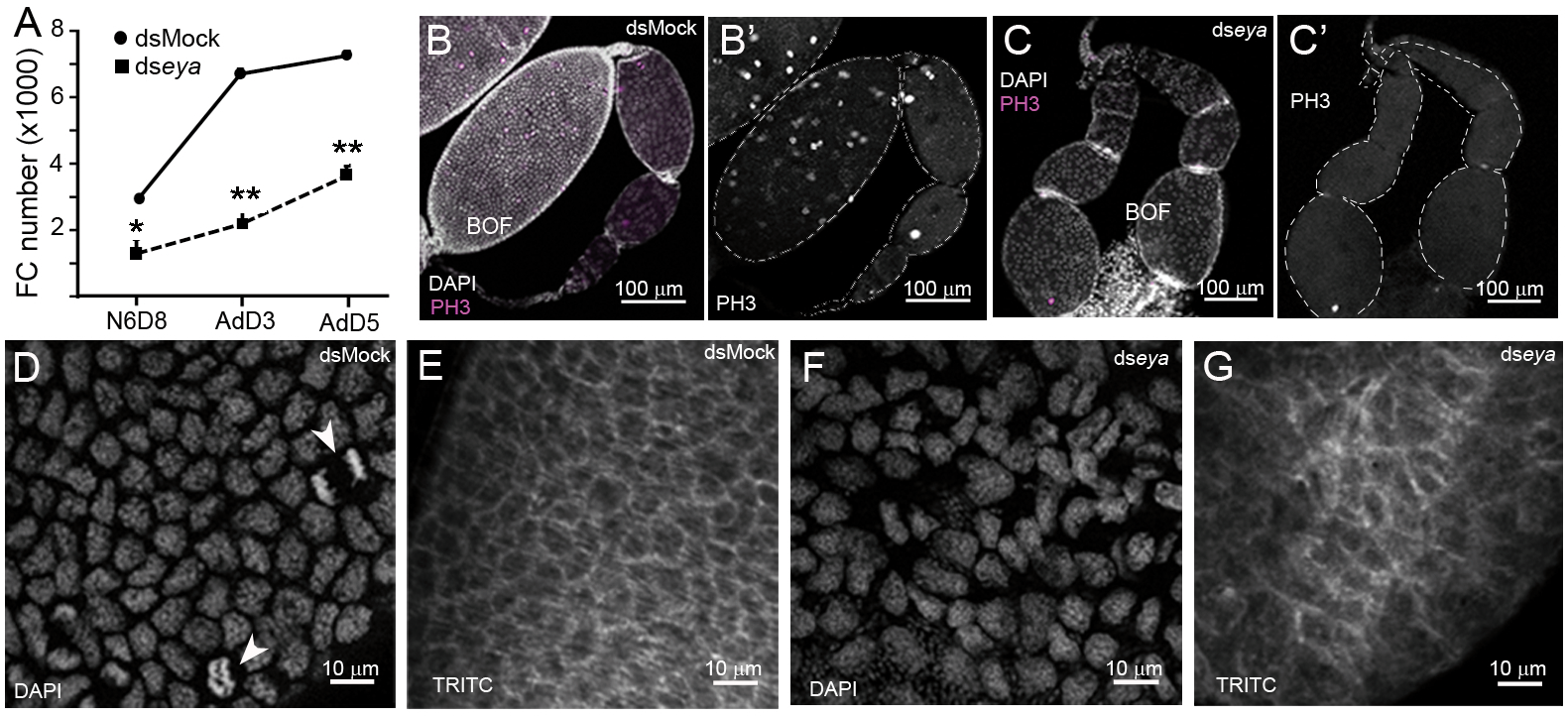
Effects of *eya* depletion in 8-day-old sixth instar nymphs of *Blattella germanica*. **A**. Number of follicular cells in basal ovarian follicles from dsMock- and *dseya*-treated females; N6D8: 8-day-old sixth instar nymph; AdD3: 3-day-old adult; AdD5: 5-day-old adult; the asterisks indicate statistically significant differences with respect to controls: *, p = 0.002 and **, p= 0.0001). **B**. Ovariole from an 8-day-old dsMock-treated nymph; the actively dividing follicular cells are labelled with anti-phospho-histone 3 (PH3) antibody (in B’, the isolated channel showing the PH3 labelling is shown; the outline of the ovarioles has been highlighted with a dashed line for clarity). **C**. Ovarioles from an 8-day-old ds*eya*-treated nymph showing a few number of cells dividing (in C’ the isolated channel showing the PH3 labelling is shown; the outline of the ovarioles has been highlighted with a dashed line for clarity); BOF: basal ovarian follicle. **D-E**. Follicular epithelia from 8-day-old dsMock-treated nymphs; In D, the follicular cell nuclei, stained with DAPI, are shown; some mitotic figures are visible (arrowheads); In E, the F-actin microfilaments, stained with TRITC, appear uniformly distributed in the cell membranes. **F-G**. Follicular epithelia from 8-day-old ds*eya*-treated nymphs. In F, the follicular cell nuclei, stained with DAPI, are shown evidencing differences in size and form with absence of mitosis; in G, the F-actin microfilaments stained with TRITC, in basal ovarian follicles display a non-uniform distribution.

These morphological changes in the ovaries became more conspicuous over time during the adult period, and basal ovarian follicles with different degrees of malformation were observed in the same ovary (Figure 3A-C). In 5-day-old adult control females, all follicular cells are binucleated and polyploid, and no further cell divisions occurs (Figure 3D and D’). F-actin are distributed on the cell membranes and appear concentrated in the expansions connecting adjacent cells (Figure 3D’’). This occurs when the follicular cells contract and leave large intercellular spaces (a phenomenon called patency, see Davey and Huebner, 1974), thus allowing the vitellogenic proteins to reach the oocyte membrane, and to be internalized into the oocyte through a specific receptor. Conversely, in 5-day-old *eya*-depleted adult females, only a few follicular cells were binucleated (Figure 3E, arrowheads), thus indicating that they became desynchronised. Besides, these cells showed high variability in size and shape, F-actins appeared concentrated in cell membranes (Figure 3E’’), and the follicular epithelia never showed signs of patency.

**Figure 3.**
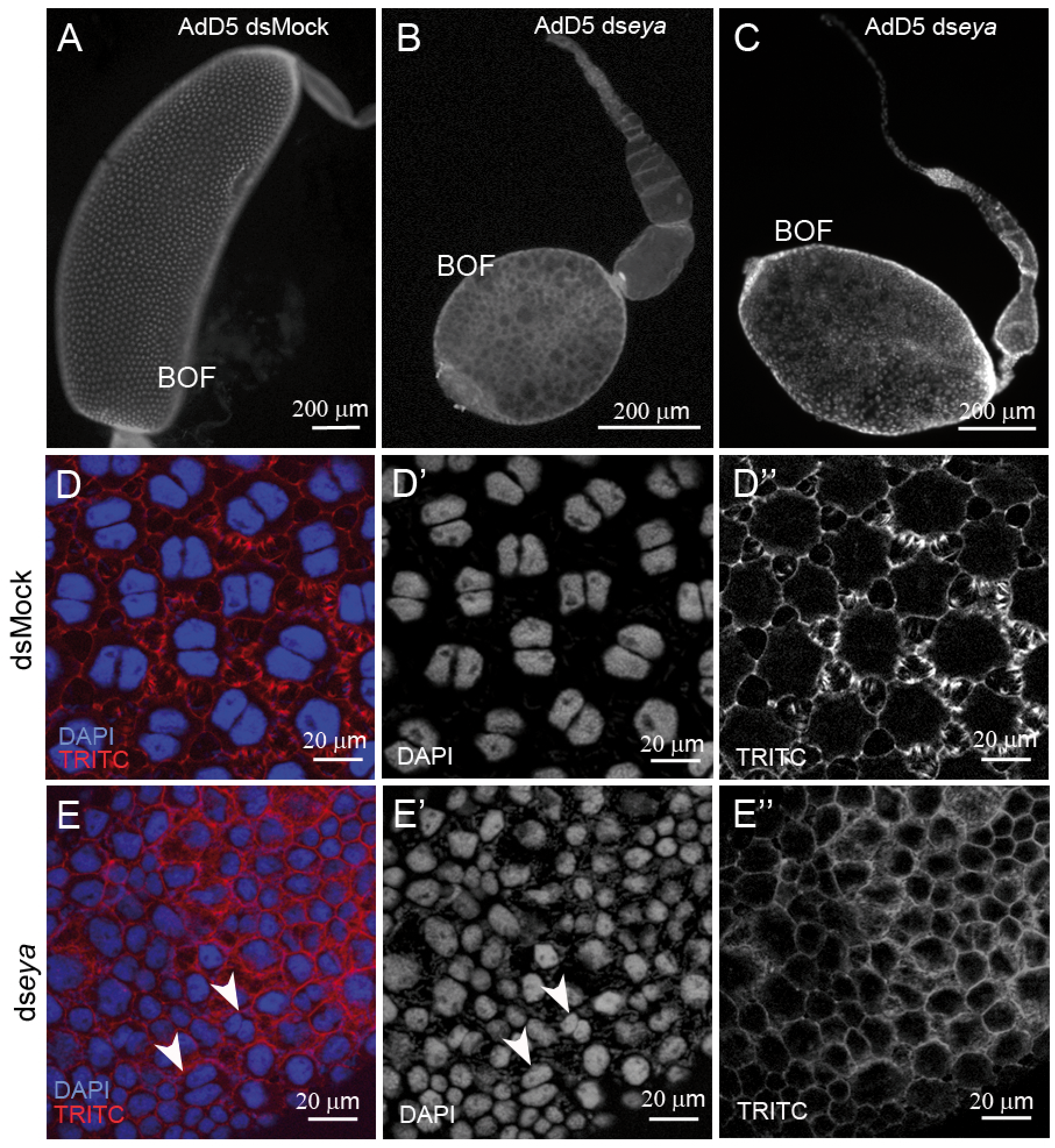
Effects of *eya* depletion in 5-day-old adult ovary of *Blattella germanica*. **A**. Ovariole from a dsMock-treated female. **B-C**. Ovarioles from ds*eya*-treated females showing different degrees of malformation; BOF: Basal ovarian follicle. **D**. Follicular epithelia of dsMock-treated females showing the binucleated cells and a high degree of patency; D’ shows the nuclei and D’’ the cytoskeleton of F-actin. **E**. Follicular epithelia of ds*eya*-treated females showing cells of different size and morphology, mostly mononucleated; E’ shows the nuclei stained with DAPI, and E’’ the cytoskeleton of F-actin, showing a uniform distribution on cell membranes and no signs of patency; arrowheads indicate a few binucleated cells.

Furthermore, the expression of the *vitellogenin receptor* (*VgR*) was upregulated in ovaries from *eya*-depleted females at the three ages examined: the last day of last nymphal instar (8-day-old) and in 3- and 5-day-old adults (p = 0.012, p = 0.022, and p = 0.001 respectively; Figure S1). Usually, *VgR* mRNA levels are high in the ovaries of newly emerged last-instar nymphs. The expression subsequently decreases until reaching very low levels in adult females at the end of vitellogenesis (Ciudad et al., 2006). The modification of basal ovarian follicle shape, together with the phenotypes observed in follicular cells and the unexpected increase in *VgR* expression in 5-day-old adult ovaries, indicate that although these ovarian follicles seemed ready to mature, they did not grow. Instead, we presumed that these ovaries might be degenerating through a programmed cell death process. To test this conjecture, we measured the expression of the effector *caspase-1* (*casp1*) in the ovaries of *eya*-depleted females. In 6- and 8-day-old sixth instar nymphs, the mRNA levels of ovarian *casp1* in *eya*-depleted insects were similar to those measured in controls (Figure S2A). However, *casp1* expression was significantly upregulated in the ovaries of 5-day-old adult *eya*-depleted insects with respect to controls (Figure S2A). In addition, labelling for the executioner *casp3* in basal ovarian follicles of 5-day-old *eya*-depleted insects appeared concentrated in the nucleus of follicular cells (Figure S2B and C), while in controls *casp3* labelling is very faint and mainly distributed throughout the cytoplasm of follicular cells. Taken together, the data suggest that basal ovarian follicles of *eya*-depleted insects are compromised at this developmental stage.

### 3.3. *eya* depletion affects somatic and germinal cells, increasing the rate of ovarian follicle differentiation

In *B. germanica* females, the number of ovarian follicles in the vitellarium is established early in the last nymphal instar, and this number is maintained during the remainder of the first gonadotrophic cycle (Table S3 and Figure 4A-C). After oviposition, a new ovarian follicle is released from the germarium to the vitellarium. This suggests that specific mechanisms maintain the number of differentiated ovarian follicles in *B. germanica*.

**Figure 4.**
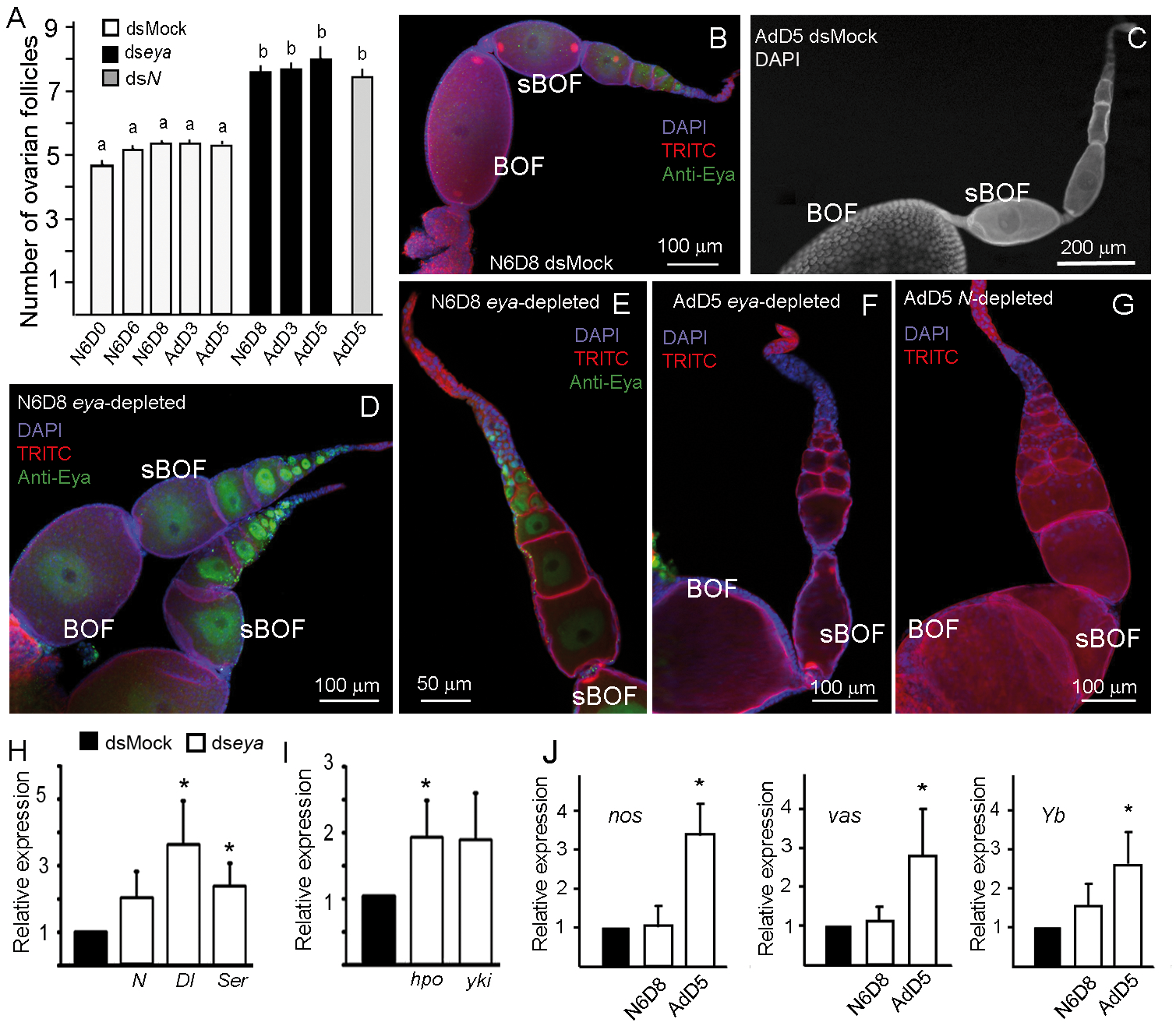
Effects of *eya* depletion on ovarian follicle differentiation in *Blattella germanica*. **A**. Number of ovarian follicles in the vitellaria from dsMock- and ds*eya*-treated females; ovarian follicles were quantified including the subbasal and all released follicles from the germarium; data from 5-day-old *N*-depleted ovarioles is also shown (obtained from Irles et al., 2016 and Irles and Piulachs, 2014); data is expressed as the mean ± S.E.M (n = 13-50) (see also Table S3); different letters indicate statistically significant differences with respect to controls (p< 0.0001). N6D0, N6D6 and N6D8: 0-day-old, 6-day-old and: 8-day-old sixth instar nymphs respectively; AdD3 and AdD5: 3-day-old and 5-day-old adult female, respectively. **B**. Ovariole from a N6D8 dsMock-treated female; BOF: basal ovarian follicle; sBOF: subbasal ovarian follicle. **C**. Vitellarium and germarium of a dsMock-treated AdD5 ovariole. **D**. Ovarioles from N6D8 ds*eya*-treated female. **E**. Vitellarium and germarium from an N6d8 ds*eya*-treated female. **F**. Ovariole from an AdD5 ds*eya*-treated female. **G**. Ovariole from AdD5; ds*N* was applied on N6D8 (see Irles et al., 2016); from B to G, the nuclei from follicular cells were stained with DAPI, F-actin microfilaments with TRICT-Phalloidin and the nucleus from germinal cells were labelled with eya 10H6 antibody (Anti-Eya). **H**. Expression of the main components of Notch pathway in ovaries from 5-day-old ds*eya*-treated adults. **I**. Expression of *hpo* and *yki* in ovaries from 5-day-old ds*eya*-treated adults. **J**. Expression of *nos, vas* and *Yb* in ovaries from 8-day-old ds*eya*-treated nymphs and 5-day-old treated adults; from H to J data represent normalized values against dsMock (reference value 1), and are expressed as the mean ± S.E.M. (n = 3); the asterisk indicates statistical significant differences with respect to controls (p< 0.02).

*eya* depletion resulted in changes in the germarium and, as a consequence, also in the vitellarium. Compared to control females, at least two extra ovarian follicles were released into the vitellarium in ovaries from 8-day-old *eya*-depleted sixth instar nymphs (Figure 4A and D-E; Table S3). These extra ovarian follicles were maintained in adult females and they were observed even in 5-day-old *eya*-depleted adult females (Figure 4A and F). In a few *eya*-depleted adult females, the vitellarium of some ovarioles contained as many as ten ovarian follicles. This concurs with the phenotype observed in *Notch* (*N*)-depleted adult females (Figure 4A and G), which is not surprising, as *N* depletion reduces *eya* expression (Irles et al., 2016; Irles and Piulachs, 2014).

In contrast, *N* expression is not affected in the ovaries of 5-day-old *eya* depleted insects, although *Delta* (*Dl*) and *Serrate* (*Ser*), the main ligands of N, appear upregulated (Figure 4H). In addition, the expression of *hippo* (*hpo*) and *yorkie* (*yki*), two important components of the Hippo pathway, significantly increases in the ovaries of *eya*-depleted females (Figure 4I). Moreover, expression of *nanos* (*nos*), *vasa* (*vas*), and *fs(1)Yb* (*Yb*), which are crucial in the modulation of germinal and somatic stem cell proliferation in *D. melanogaster* (King et al., 2001; Wang and Lin, 2004), results significantly upregulated in the ovaries of 5-day-old *B. germanica eya*-depleted insects (Figure 4I). Taken together, the results indicate that *eya*-depletion affects differently the distinct regions of the ovary. In basal ovarian follicles, their development becomes arrested, and they tend to die (Figure S2), whereas in the germarium, *eya*-depletion triggers an increase in differentiated cells, thus raising the number of ovarian follicles in the vitellarium (Figure 4).

### 3.4. Ecdysone signalling and the differentiation of ovarian follicles

In *D. melanogaster*, the formation and differentiation of ovarian follicles are triggered by 20E signalling (see Ameku et al., 2017; Belles and Piulachs, 2015; Hsu et al., 2019; König et al., 2011; Uryu et al., 2015). Therefore, we presumed that this mechanism might also operate in *B. germanica*. The prothoracic gland is the main source of ecdysteroids in *B. germanica* nymphs whereas the ovary is its main source in the adult female, as the prothoracic gland degenerates after metamorphosis (Belles, 2020; Pascual et al., 1992; Romaña et al., 1995). However, it has not been described wether *B. germanica* nymphal ovaries can produce ecdysone/20E, and/or respond to it, and regulate developmental processes.

To examine the response of nymphal ovaries to an ecdysone/20E signal, the expression of one ecdysone-dependent early gene, *E75A* (Mané-Padrós et al., 2008), was measured in ovaries of *B. germanica* during the sixth nymphal instar and the adult.The expression pattern of *E75A* correlates with the profile of ecdysteroids titre in the haemolymph in last nymphal instar, and with the levels of ecdysteroids in the adult ovary, where the peak of ecdysone during choriogenesis coincides with the maximal expression of *E75A* (Figure 5A).

**Figure 5.**
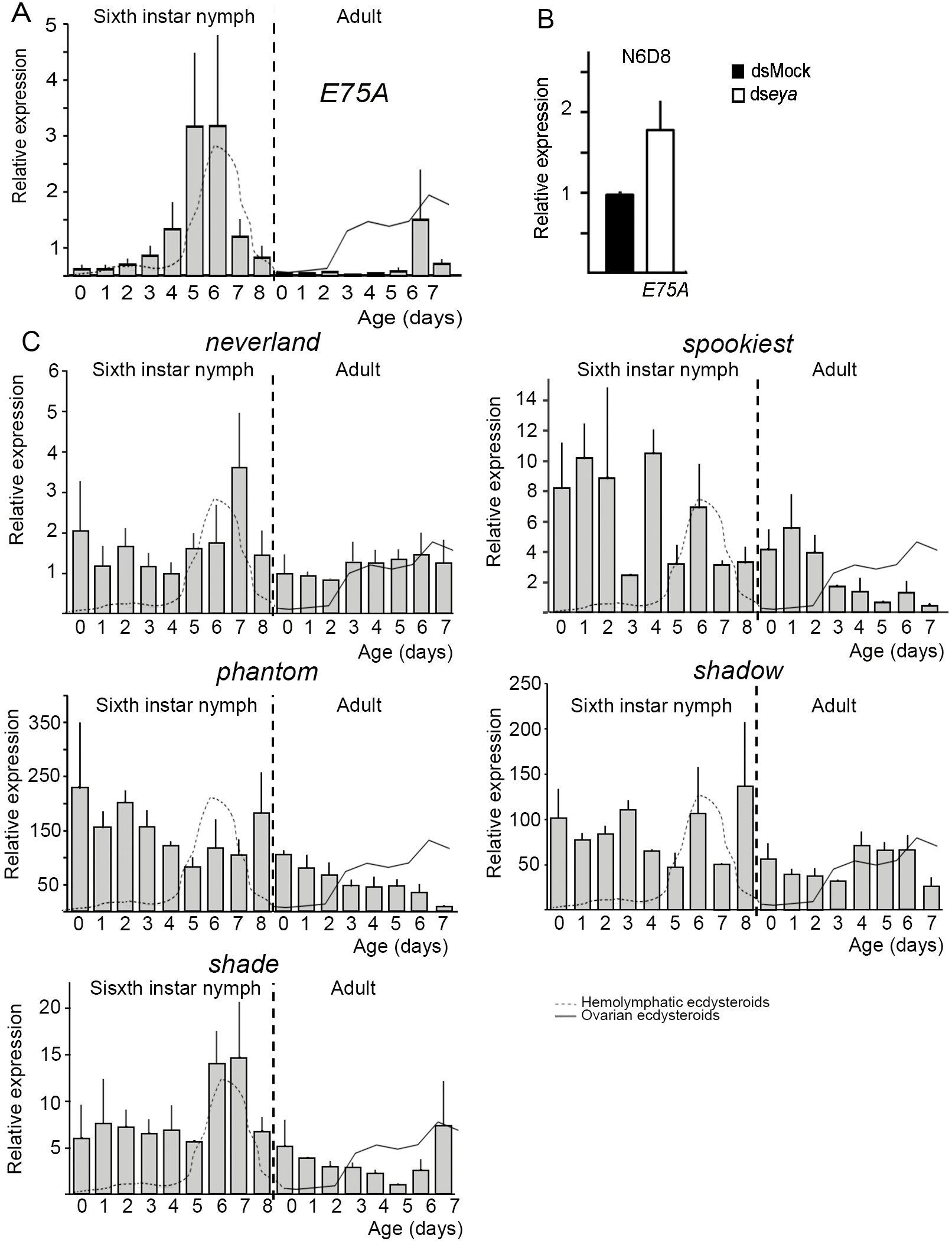
Expression of *E75A* and steroidogenic genes in the ovaries of *Blattella germanica*. **A**. The expression pattern of *E75A* in ovaries of sixth instar nymphs and adult females. **B**. Expression of *E75A* in ovaries from 8-day-old ds*eya*-treated nymphs (N6D8); data represent normalized values against dsMock (reference value 1), and are expressed as the mean ± S.E.M. (n = 3). **C**. The mRNA expression patterns of *neverland, spookiest, phantom, shadow* and *shade*, in ovaries of sixth instar nymphs and adult females; in A and C, the profiles of ecdysteroid titer in the haemolymph (grey dashed line), and ecdysteroid content in the ovaries (grey solid line) are also shown (data from Cruz et al., 2003; Pascual et al., 1992; Romaña et al., 1995); data represent copies of mRNA per 1000 copies of *actin-5c* (relative expression) and are expressed as the mean ± S.E.M. (n= 3-6).

Furthermore, the expression of *E75A* was measured in ovaries from 8-day-old *eya*-depleted last instar nymphs. Results showed that *E75A* expression was upregulated on average, although the differences between controls and ds*eya*-treated were not statistically significant (Figure 5B). This suggests that *eya*-depletion triggered an increase of ecdysone signalling in the nymphal ovaries.

Although the *B. germanica* adult ovary is able to produce ecdysteroids (Cruz et al., 2003; Pascual et al., 1992), the expression of steroidogenic genes in the adult ovary has not been explored. Hence, we measured the expression of *neverland* (*nvd*), *spookiest* (*spot*), *phantom* (*phm*), *shadow* (*sad*) and *shade* (*shd*), during the sixth nymphal instar and the adult. The results (Figure 5C) showed that the expression patterns of the different genes do not correlate with each other, or with the profiles of ecdysteroids. However, the expression levels are higher in the last nymphal instar than in the adult, in general. Moreover, the highest expression levels of *nvd* and *shd* appear to coincide with the maximum peak of ecdysteroids, in the last nymphal instar, and with the highest content of ecdysteroids in the adult ovary (Figure 5C).

Furthermore, in the ovaries of 8-day-old *eya*-depleted last instar nymphs, we measured the expression of *nvd, spot* and *shd*, three genes that represent three characteristic steps of the biosynthesis of 20E, an early step (*nvd*), a step in the so called black box (*spot*) and that of the transformation of ecdysone into 20E (*shd*) (Niwa and Niwa, 2016; Ou et al., 2016). Results indicated that the expression of *nvd* and s*hd* was not significantly affected by *eya* depletion, whereas that of *spot* was upregulated (Figure 6A). These results suggest that *eya* represses the expression of at least *spot*, which might compromise the ecdysone biosynthesis in the ovaries, thus possibly affecting oogenesis processes.

**Figure 6.**
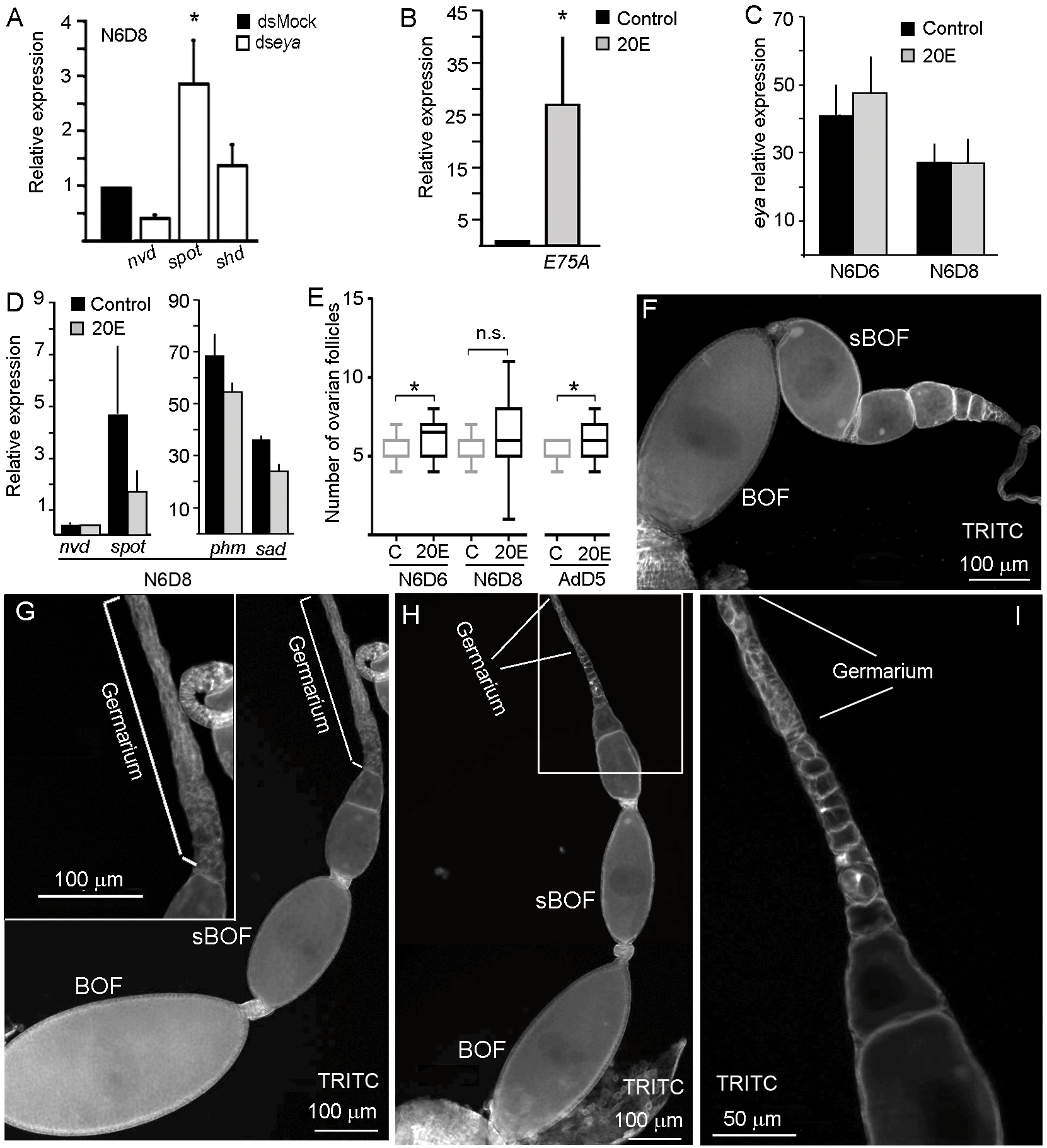
Effects of 20E treatment on ovarian development in *Blattella germanica*. **A**. Expression levels of *nvd*, s*pot* and s*hd*, in ovaries from N6D8 ds*eya*-treated females; data represent normalized values against the control (reference value 1) and are expressed as the mean ± S.E.M. (n = 4 −10); the asterisk indicates statistical significant differences with respect to controls (p <0.002). **B**. Expression levels of *E75A* in ovaries from N6D8 treated with 20E; data represent normalized values against the control (reference value 1) and are expressed as the mean ± S.E.M. (n = 4 - 10); the asterisk indicates statistical significant differences with respect to controls (p <0.001). **C**. Expression of *eya* in nymphal ovaries from females treated with 20E. **D**. Expression levels of n*vd*, s*pot, phm* and s*ad* in ovaries from N6D8 treated with 20E; data represent copies of mRNA per 1000 copies of a*ctin-5c* and are expressed as the mean ± S.E.M. (n = 3-4). **E**. Box plot representing the number of ovarian follicles localized in the vitellarium and germarium in 20E-treated females, the asterisk indicates statistical significant differences with respect to controls (p < 0.002; n.s. no significant;n= 20-50). **F**. Ovariole from a control N6D8; BOF: Basal ovarian follicle, sBOF: subbasal ovarian follicle. **G**. Ovariole from a N6D8 20E-treated female, showing a reduced number of ovarian follicles released from the germarium; in the inset, the germarium is show at higher magnification. **H**. Ovariole from a N6D8 20E-treated female, showing a high number of ovarian follicles released from the germarium. **I**. Detail of the germarium from panel H, at higher magnification showing the differentiated ovarian follicle; samples were stained with phalloidin-TRITC to show the F-actin microfilaments. From B to I, last instar nymphs and adult females were treated with 10 µM 20E at the day of emergence to the respective instar, whereas controls were treated with a solution of 10% EtOH; N6D6: 6-day-old sixth instar nymph; N6D8: 8-day-old sixth instar nymph; AdD5: 5-day-old adult female.

To test the effects of ecdysteroids on *E75A* expression in the ovaries, we applied 20E to newly emerged last instar nymphs. Then, we measured the expression of *E75A* in the ovaries of 8-day-old last instar nymphs. Results showed that *E75A* expression was significantly upregulated in the ovaries of 20E-treated insects, with a fold change close to 30 (Figure 6B). This indicates that *E75A* readily respond to 20E in nymphal ovaries, which suggest that its expression can be used as readout of ecdysteroid changes, as used in other studies (Colombani et al., 2012; Li et al., 2016).

Additionally, results showed that *eya* expression in the ovaries of the 8-day-old last instar nymphs was not affected by the 20E treatment (Figure 6C). Similarly, the expression of the steroidogenic genes did not significantly change after 20E treatment (Figure 6D). Regarding the ovarian follicles in the vitellarium, its number increased after the 20E treatment (Figure 6E; Table S3). In 6-day-old last instar nymphs, a significant increase (p < 0.002) in the number of differentiated ovarian follicles was observed. However, two days later, the number of ovarian follicles found in the vitellarium was very variable (ranging between 3 and 11; Figure 6F and G-H’) compared to the controls (Figures 6E and F), which suggests that some ovarian follicles underwent cell death. Finally, the effect of 20E on ovarian follicle differentiation was not limited to the last nymphal instar. Newly emerged adult females that had been treated with 20E also showed a higher number of differentiated ovarian follicles in comparison with the respective controls (p< 0.002; Figure 6E).

## 4. Discussion

In hemimetabolan species, *eya* was originally described by its involvement in eye development, which was related to cell proliferation (Dong and Friedrich, 2010; Takagi et al., 2012). In the present work we have shown the involvement of *eya* in ovary development in a hemimetabolan species. Depletion of *eya* in the cockroach *B. germanica* prevents the completion of the gonadotrophic cycle, which derives in females sterility.

The phenotypes observed after *eya* depletion in last instar nymphs of *B. germanica* indicate that this gene acts early in oogenesis, playing distinct roles in different regions of the panoistic ovary. Indeed, the development of basal ovarian follicles, which usually start to grow and mature during the last nymphal instar, is arrested in *eya*-depleted females. Moreover, the basal ovarian follicles lost their typical elliptical morphology and become spherical. A similar phenotype was observed in *N*-depleted females of *B. germanica* (Irles and Piulachs, 2014), and, intriguingly, this phenotype becomes more conspicuous as the females aged. The fact that *eya* depletion partially phenocopies N-depletion is no surprising, as *N* depletion results in a decrease of *eya* expression (Irles and Piulachs, 2014). In contrast, the disappearance of the stalks between the ovarian follicles in the ovariole, which is an additional consequence of *N* depletion (Irles and Piulachs, 2014), was not observed in *eya*-depleted insects, whose ovaries show a well formed stalk between the basal and sub-basal ovarian follicles, although the stalk between the youngest ovarian follicles was frequently absent or undifferentiated.

The fat body of eya-depleted females is expressing vitellogenin at similar levels than in controls (results not shown). However, the observations in *eya*-depleted females suggest that vitellogenin is not incorporated into the growing oocytes, which might be due to the absence of the corresponding receptor, VgR. Intriguingly, abundant *VgR* transcripts accumulate in ovaries of *eya*-depleted females, whereas the expression of *VgR* mRNA levels decreases in control females as the oocyte growth, a decrease that coincides with the increase of VgR protein levels in the membrane of basal oocytes (Ciudad et al., 2006). Taken together, the data suggest that *VgR* translation is prevented in *eya*-depleted females, explaining why the basal oocytes do not incorporate vitellogenin.

The most remarkable phenotype observed in ovarioles from *eya*-depleted females was the uncontrolled cell proliferation and differentiation in the germaria, resulting in an increase in the number of differentiated ovarian follicles produced. Interestingly, they show a notable swelling of the germaria, a phenotype that is reminiscent of that described in *D. melanogaster eya*-null mutants (Bai and Montell, 2002; Bonini et al., 1998). The aforementioned swollen shape occurred in these *eya-*null mutant flies because the development of the maturing egg chamber arrested, but the germaria continued proliferating (Bonini et al., 1998). This concurrence suggests that the role of *eya* in the control of stem cell differentiation and proliferation has been evolutionarily conserved between cockroaches and flies.

In *D. melanogaster*, the formation and differentiation of ovarian follicles is triggered by 20E (see Ameku et al., 2017; Belles and Piulachs, 2015; Hsu et al., 2019; König et al., 2011; Uryu et al., 2015). The triggering effect of 20E on somatic and germinal cells in the germarium, appear ancestral and also operating in less modified insects, where juvenile hormone plays a gonadotropic role (Belles et al., 2000; Comas et al., 2001; Treiblmayr et al., 2006).The source of ecdysone in adult *B. germanica* females is the ovary, and at this age, their main function is to promote chorion synthesis in mature basal ovarian follicles (Pascual et al., 1992; Romaña et al., 1995). However, steroidogenic genes are expressed in ovaries of last instar nymphs of *B. germanica*, which suggest that immature ovaries of this species also synthesise ecdysone.

In the germarium of *D. melanogaster*, ecdysone signalling controls de quantity, but not the differentiation status of germinal stem cells. In fact, the latter may be mediated by the Notch pathway. Ecdysone signalling induces *Dl* expression at the terminal filament in cell membranes, which activates *N* and determines the fate of these cells, which can become cap or escort cells (Ameku et al., 2017; Green et al., 2011; Hsu et al., 2019). Our results in *B. germanica* suggest that similar signalling networks can occur in panoistic ovaries. When *eya* is depleted *Dl* expression significantly increase and *N* expression becomes activated, again suggesting that ecdysone levels had increased. Thus, the signalling pathway found in *B. germanica* appears equivalent to that described in *D. melanogaster*, and, significantly, *eya*-depletion in cockroaches results in swollen germaria in the ovarioles, as occurs in the fruit fly.

The overexpression of steroidogenic genes in the ovaries of *eya*-depleted nymphs suggests that *eya* regulates the proliferation of ovarian follicles in the *B. germanica* ovary by controlling the steroidogenic pathway. This idea is in line with the expression of *eya* in adult ovaries, in which the decrease of *eya* levels at the end of the gonadotrophic cycle coincides with the increase of ovarian ecdysone at this age (Romaña et al., 1995). This ecdysone increase triggers the production of chorion proteins in mature basal ovarian follicles (Pascual et al., 1992).

Ectopic treatment with 20E gave similar results to those obtained after *eya* depletion: ovarian follicle proliferation, and swelling of the germaria. However, *eya* expression was not modified by 20E treatment, which indicates that *eya* regulates the activity of the steroidogenic pathway but is not controlled by 20E.

In summary in *B. germanica, eya* as a downstream component of the Notch pathway, regulates the correct cell proliferation and cell fate in the follicular epithelia of basal ovarian follicles. While in the germarium and the terminal filament, *eya* acts on somatic and germinal cells regulating their differentiation and proliferation, by controlling ecdysone signalling. Important from an evolutionary point of view, these functions are equivalent, possibly homologous, to the functional duality of *eya* reported in the most derived fly *D. melanogaster*.

## Acknowledgments

We thank the financial support from the Ministry of Science and Innovation, Spain for grants BFU2011-22404 and CGL2016-76011-R that also received financial assistance from the European Fund Economic and Regional Development (FEDER) And from the Catalan Regional Government (grant 2017 SGR 1030). PI is received the financial support from *Apoyo al Retorno de Investigadores desde el Extranjero* program (*Convocatoria* 2013, 821320046, PAI, CONICYT). The funders had no role in the design of the study, data collection or analysis, the decision to publish, or in the preparation of the manuscript.

## Figure Legends

**Figure S1.**
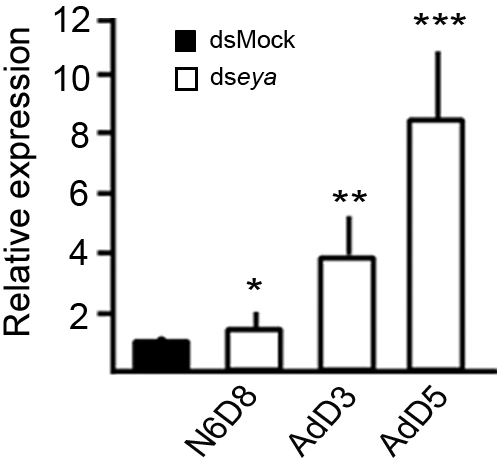
Expression of *Vitellogenin receptor* in ovaries from eya-depleted females. Newly emerged last instar nymphs were treated with dseya and ovaries dissected at different ages. N6O8: 8-day-old sixth instar nymph; AdO3: 3-day-old adult; Ad 05: 5-day-old adult. Data represent normalized values against the control (reference value 1) and are expressed as the mean ± S.E.M. (n = 3). The asterisks indicate statistically significant differences (t-test) with respect to controls:*, p = 0.012; **, p= 0.022; ***, p = 0.001.

**Figure S2.**
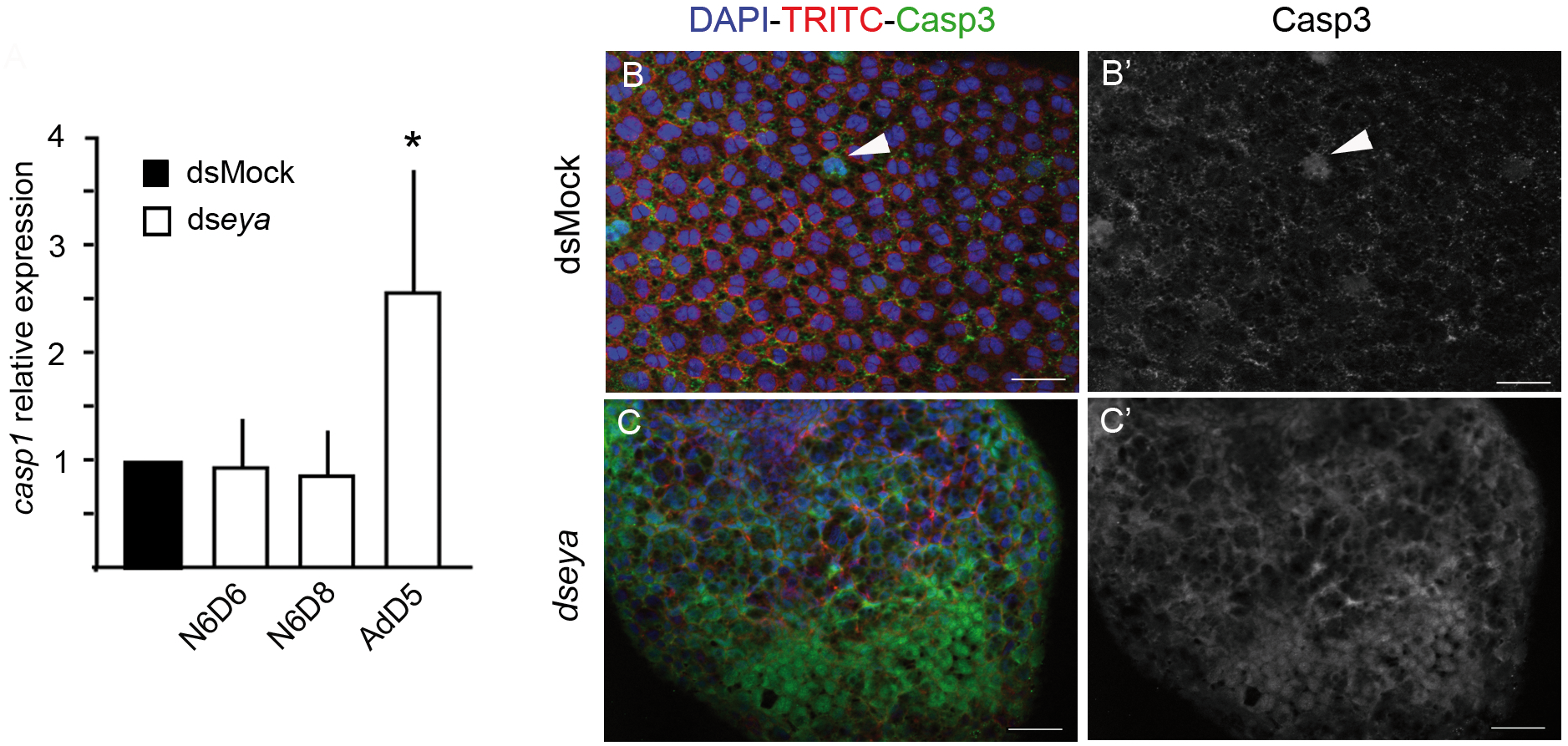
Caspase activity in eya-depleted ovaries. **A**. Expression of *caspase 1 (casp1)* in ovaries from eya-depleted females. Newly emerged last instar nymphs were treated with dseya RNA, and ovaries dissected at different ages. N6D6: 6-day-old sixth instar female; N6D8: 8-day-old sixth instar female; Ad 05: 5-day-old adult female. Data represent normalized values against the control (reference value 1) and are expressed as the mean± S.E.M. (n = 3). The asterisks indicate statistically significant differences (t-test) with respect to controls: p < 0.001. **B**. Follicular epithelium in basal ovarian follicle from a 5-day-old dsMock-treated female. The uniform distribution of binucleated follicular cells is showed. Only some sporadic nuclei (arrowhead) appeared labelled by the anti-Casp3 antibody (green). In **B’** the labelling for Casp3 is shown. **C**. Follicular epithelium in basal ovarian follicle from a 5-day-old eya-depleted female, most of the nuclei appeared labelled for Casp3. In C’ nuclei labelled for Casp3 are shown. In all images, the posterior pole of the basal follicle is towards the bottom. Scale bars: 50 µm.

**Table S1:**
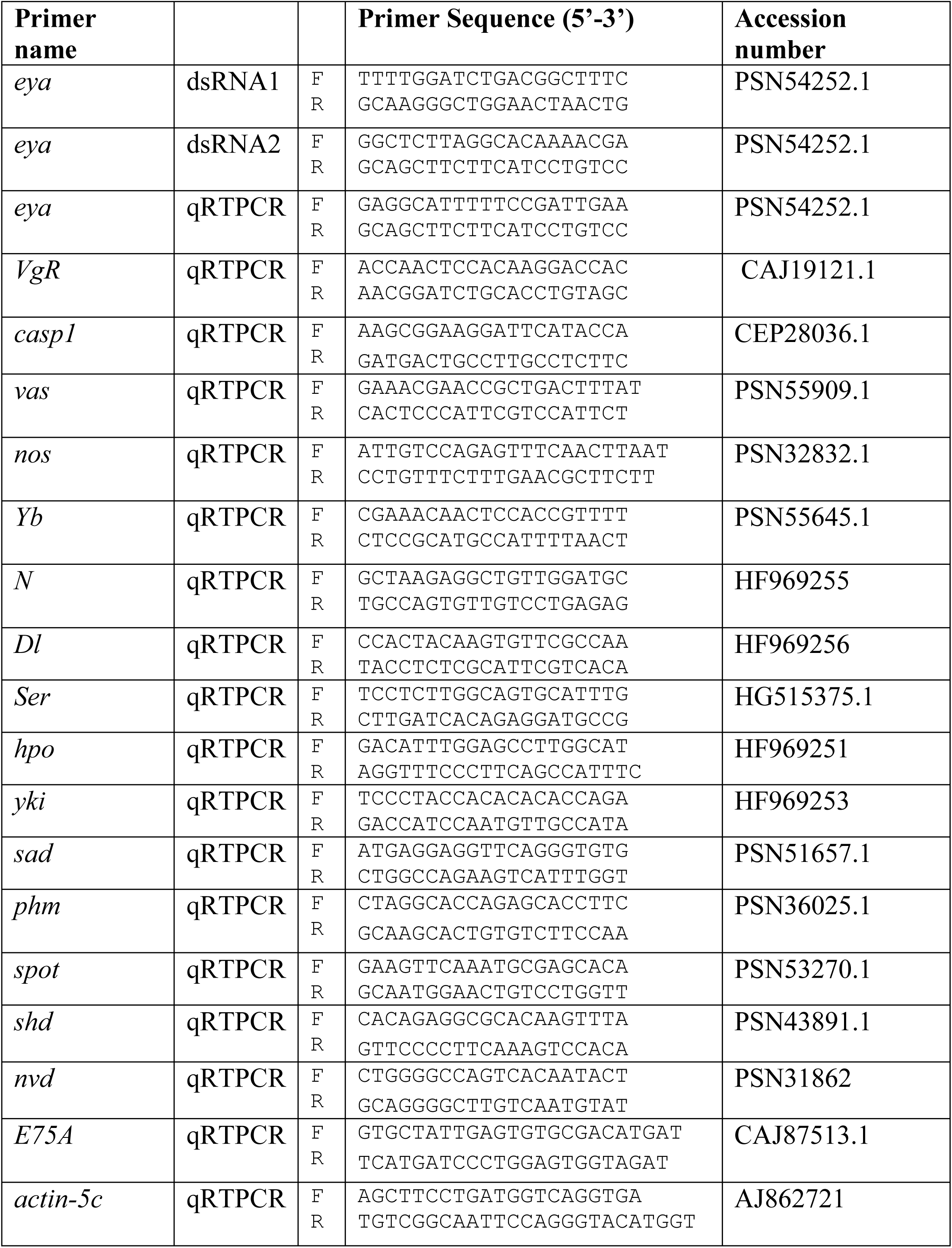
Primer sequences used for quantitative real-time PCR (qRT-PCR) and RNAi experiments. Actin-5c is the housekeeping gene use in expression studies. F: Primer forward; R: Primer reverse

**Table S2.** *eya* depletion in oviposition. Newly emerged sixth instar nymphs (N6D0) females were treated with ds*eya* or dsMock and left to oviposit. The day of oviposition (from the day of adult emergence), the number of females that oviposit, the days that the oothecae were transported and the number of fertile oothecae were recorded.

| Treatment (day of treatment) | n | Days to oviposit | % Females that oviposit | Days of oothecae transport | % Fertil oothecae |
| --- | --- | --- | --- | --- | --- |
| *dsMock* (N6D0) | 40 | 7.68 ± 0.13 | 100 | 17.92 ± 0.11 | 100 |
| *dseya* (N6D0) | 36 | - | 0 | - | - |

**Table S3.** Number of ovarian follicles (OF) in the vitellaria from dsMock-, ds*eya*- and 20E treated females, were quantified at different ages. Data was recorded from different ovarioles (n) belonging to 7-10 females. Considering those OF from the subbasal position to the youngest that has left the germarium, both included.

| Treatment | Age | n | OF (mean ± SEM) |
| --- | --- | --- | --- |
| **dsMock (N6D0)** | N6D0 | 22 | 4.68 ± 0.166 |
| | N6D6 | 50 | 5.48 ± 0.131 |
| | N6D8 | 30 | 5.37 ± 0.122 |
| | AdD3 | 31 | 5.32 ± 0.169 |
| | AdD5 | 28 | 5.32 ± 0.126 |
| **dseya (N6D0)** | N6D8 | 50 | 7.60 ± 0.191 |
| | AdD3 | 13 | 7.69 ± 0.208 |
| | AdD5 | 13 | 8.00 ± 0.408 |
| **20E (N6D0)** | N6D6 | 18 | 6.28 ± 0.289 |
| | N6D8 | 39 | 6.10 ± 0.319 |
| **20E (AdD0)** | AdD5 | 28 | 6.20 ± 0.205 |

